# Taxonomic classification methods reveal a new subgenus in the paramyxovirus subfamily *Orthoparamyxovirinae*

**DOI:** 10.1101/2021.10.12.464153

**Authors:** Heather L. Wells, Elizabeth Loh, Alessandra Nava, Mei Ho Lee, Jimmy Lee, Jum R. A. Sukor, Isamara Navarrete-Macias, Eliza Liang, Cadhla Firth, Jonathan Epstein, Melinda Rostal, Carlos Zambrana-Torrelio, Kris Murray, Peter Daszak, Tracey Goldstein, Jonna A.K. Mazet, Benhur Lee, Tom Hughes, Edison Durigon, Simon J. Anthony

**Affiliations:** Department of Ecology, Evolution, and Environmental Biology, Columbia University, New York, NY, USA; EcoHealth Alliance, New York, NY, USA; Instituto Leônidas & Maria Deane, Fiocruz Amazônia, Manaus, BR; Conservation Medicine, Selangor, Malaysia; Sabah Wildlife Department, Kota Kinabalu, Sabah, Malaysia; Department of Pathology, Microbiology, and Immunology, School of Veterinary Medicine, University of California Davis, Davis, CA, USA; Center for Infection and Immunity, Mailman School of Public Health, Columbia University, New York, NY, USA; School of Public Health, Imperial College London, London, UK; MRC Unit The Gambia at London Schol of Hygiene and Tropical Medicine, Atlantic Boulevard, Fajara, The Gambia; Emerging Threats Division, Office of Infectious Disease, Bureau for Global Health, U.S. Agency for International Development, Washington D.C., USA; One Health Institute and Karen C. Drayer Wildlife Health Center, School of Veterinary Medicine, University of California, Davis, CA, USA; Icahn School of Medicine at Mount Sinai, New York, NY, USA; Institute of Biomedical Science, University of Sao Paulo, Sao Paulo, BR

## Abstract

As part of a broad One Health surveillance effort to detect novel viruses in wildlife and people, we report several paramyxoviruses sequenced primarily from bats during 2013 and 2014 in Brazil and Malaysia, including seven from which we recovered full-length genomes. Of these, six represent the first full-length paramyxovirus genomes sequenced from the Americas, including two sequences which are the first full-length bat morbillivirus genomes published to date. Our findings add to the vast number of viral sequences in public repositories that have been increasing considerably in recent years due to the rising accessibility of metagenomics. Taxonomic classification of these sequences in the absence of phenotypic data has been a significant challenge, particularly in the paramyxovirus subfamily *Orthoparamyxovirinae*, where the rate of discovery of novel sequences has been substantial. Using pairwise amino acid sequence classification (PASC), we describe a novel genus within this subfamily tentatively named *Jeishaanvirus*, which we propose should include as subgenera *Jeilongvirus, Shaanvirus*, and a novel South American subgenus *Cadivirus*. We also highlight inconsistencies in the classification of Tupaia virus and Mojiang virus using the same demarcation criteria and show that members of the proposed subgenus *Shaanvirus* are paraphyletic. Importantly, this study underscores the critical importance of sequence length in PASC analysis as well as the importance of biological characteristics such as genome organization in the taxonomic classification of viral sequences.

## Introduction

With the recent emergence of zoonotic paramyxoviruses, including Nipah virus and Hendra virus, a great deal of effort has been placed on discovering novel paramyxoviruses in wildlife [1– 3]. However, increased surveillance over the last decade has revealed a multitude of novel bat- and rodent-borne paramyxoviruses (PMVs) that do not fall into previously defined genera and are therefore difficult to classify. This issue has been particularly problematic within the *Orthoparamyxovirinae*, where most newly discovered sequences are phylogenetically positioned between the well-established genera *Morbillivirus* and *Henipavirus* [4]. Recently two new genera have been named to encompass these sequences (*Jeilongvirus* and *Narmovirus*) [5]; however, these new taxonomic classifications have not kept pace with the increasing rate of discovery of additional novel sequences that do not appear to fall within any of these four currently named genera (Figure 1).

**Figure 1.**
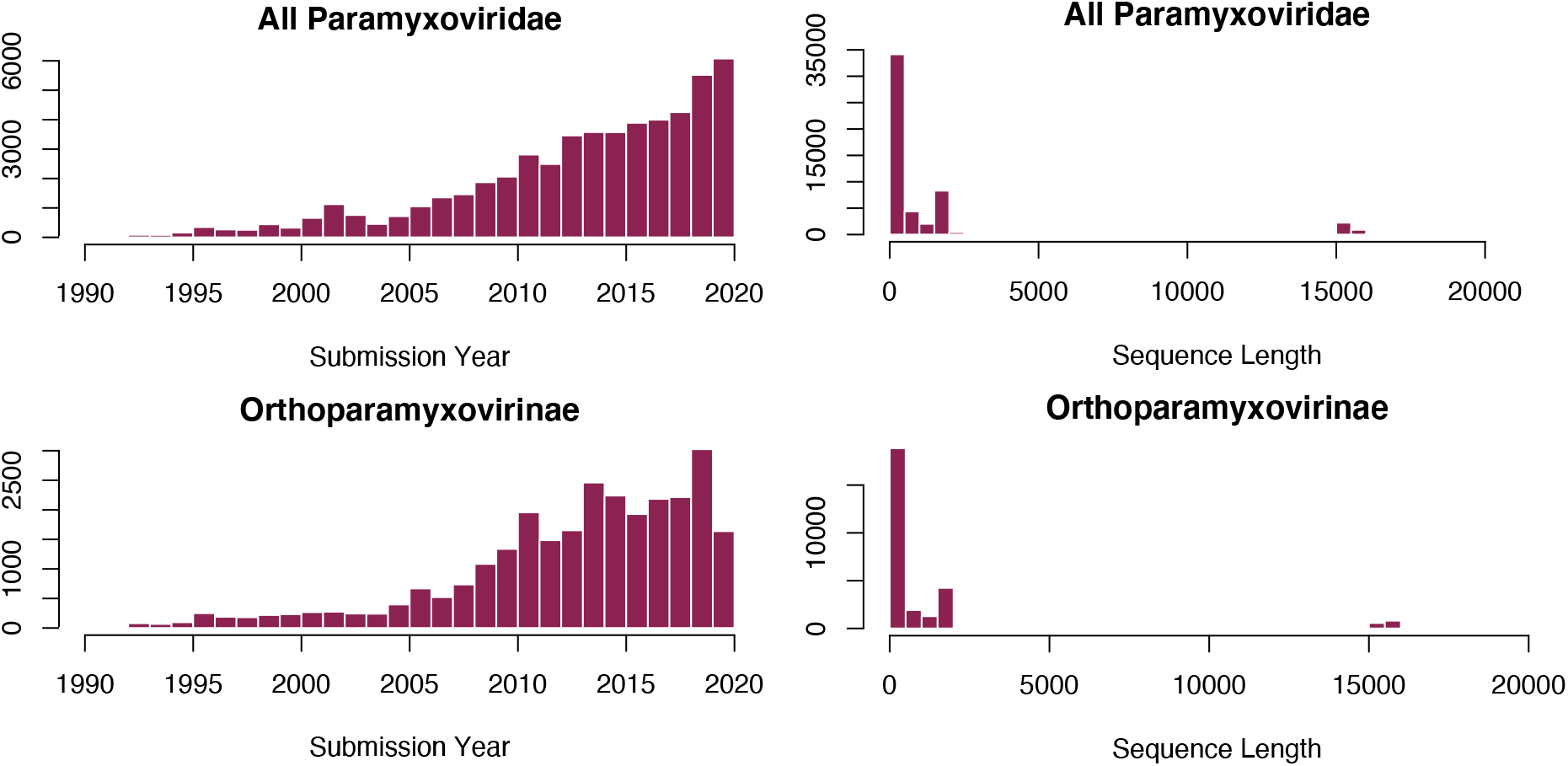
Histogram of paramyxovirus sequences submitted to GenBank by submission year (left) and sequence length (right). Trends are shown for all *Paramyxoviridae* (top) and *Orthoparamyxovirinae* only (bottom). The number of sequences submitted is increasing year over year, but the significant majority of sequences submitted are short PCR fragments (∼500bp) and very few are full genomes (∼15-16kb).

Classically, paramyxovirus taxonomy has relied on phenotypic differences such as the presence of neuraminidase and/or haemagglutination activity to demarcate distinct taxonomic groups [4,6,7]. However, the rate of detection of new viruses through genetic sequencing has now vastly outpaced the rate at which these viruses can be isolated and experimentally characterized, necessitating a new taxonomic approach. New frameworks for virus classification emphasize a wider biological context, such as host range, genome organization and presence of additional transcriptional units (ATUs), pairwise amino acid sequence comparison (PASC), and cell receptor usage [4,6]. Recently, several authors have suggested the creation of a new paramyxovirus genus, *Shaanvirus*, to encompass a group of sequences that are related to jeilongviruses but have a distinct genome organization and host range [8–10]. These new viruses have been sampled from bats (with the exception of Belerina virus, which was found in a hedgehog) as opposed to rodents, which are the only host group known to harbor jeilongviruses [11–14], and have non-homologous ORFs between the fusion protein and receptor binding protein (RBP) genes in the genome.

Here, we report the sequences of seven new paramyxovirus genomes from bats: two within the genus *Morbillivirus* from Brazil, one closely related to the proposed *Shaanvirus* group from Sabah on Malaysian Borneo, and four additional sequences from Brazil which form a monophyletic group and have biological features distinct from any currently named genera. We find that a modified version of the pairwise amino acid sequence comparison (PASC) tool does not support the classification of these four genomes as a new genus, despite the distinct characteristics of this clade. Instead, our investigations support the classification of these four novel sequences as a subgenus along with *Shaanvirus* and *Jeilongvirus* into a single genus, with *Shaanvirus* and *Jeilongvirus* also becoming subgenera. These three groups have the defining commonality of additional transcriptional units in the genome not present in other orthoparamyxoviruses. We also highlight a number of considerations for the existing classification of other orthoparamyxoviruses: (1) the proposed *Shaanvirus* group is paraphyletic and contains two groups of sequences with different biological contexts; and (2) the classification of Mojiang and Tupaia viruses are inconsistent with our proposed genus demarcation cutoffs. Critically, we also show that previous limitations to using PASC as a tool to classify sequences can be resolved when specific regard is given to the length of the sequences being tested.

## Methods

### Field sampling and laboratory screening for PMVs

Rectal, oral, blood, and urine samples from bats, rodents, or non-human primates in Brazil and Sabah, Malaysia were previously collected as part of a larger research effort [15] and analyzed for this study (Supplementary Tables 1 and 2). Extractions of total nucleic acid were performed and converted to cDNA as previously described [16,17]. Consensus PCR (cPCR) assays using *Paramyxoviridae*-specific degenerate primers were used to screen for samples positive for paramyxoviruses as described in Tong et al. [18]. The degenerate primers bind two highly conserved regions with variable sequence in between, allowing for broad reactivity within the viral family and detection of both known and novel viruses. PCR products were screened by gel electrophoresis, and bands of the expected amplicon size were cloned using Strataclone PCR cloning kits and sequenced using Sanger sequencing to confirm the presence of paramyxoviruses. Partial gene sequences of positive samples were deposited in GenBank (Supplementary Tables 1 and 2).

### Genome sequencing of PMV-positive samples

Total nucleic acid of 39 paramyxovirus-positive samples were sequenced using the Illumina HiSeq platform. Quality control and adapter trimming were performed on the resulting reads using Cutadapt v1.18 and host reads were subtracted using Bowtie v2.3. 5 (*Phyllostomus*: NCBI Genome ID 75334; *Carollia* and *Diamus*: 22833, *Hipposideros*: 75235; *Myotis*: 43810).

Resulting reads were *de novo* assembled using MEGAHIT v1.2.8 [19]. Contigs were scaffolded to a reference sequence (morbilliviruses: GenBank accession AF014953; jeishaanviruses: KC154054) and any overlaps or gaps were confirmed with iterative local alignment using Bowtie2 [20]. The full genome sequences are deposited in GenBank (Table 1).

**Table 1.**
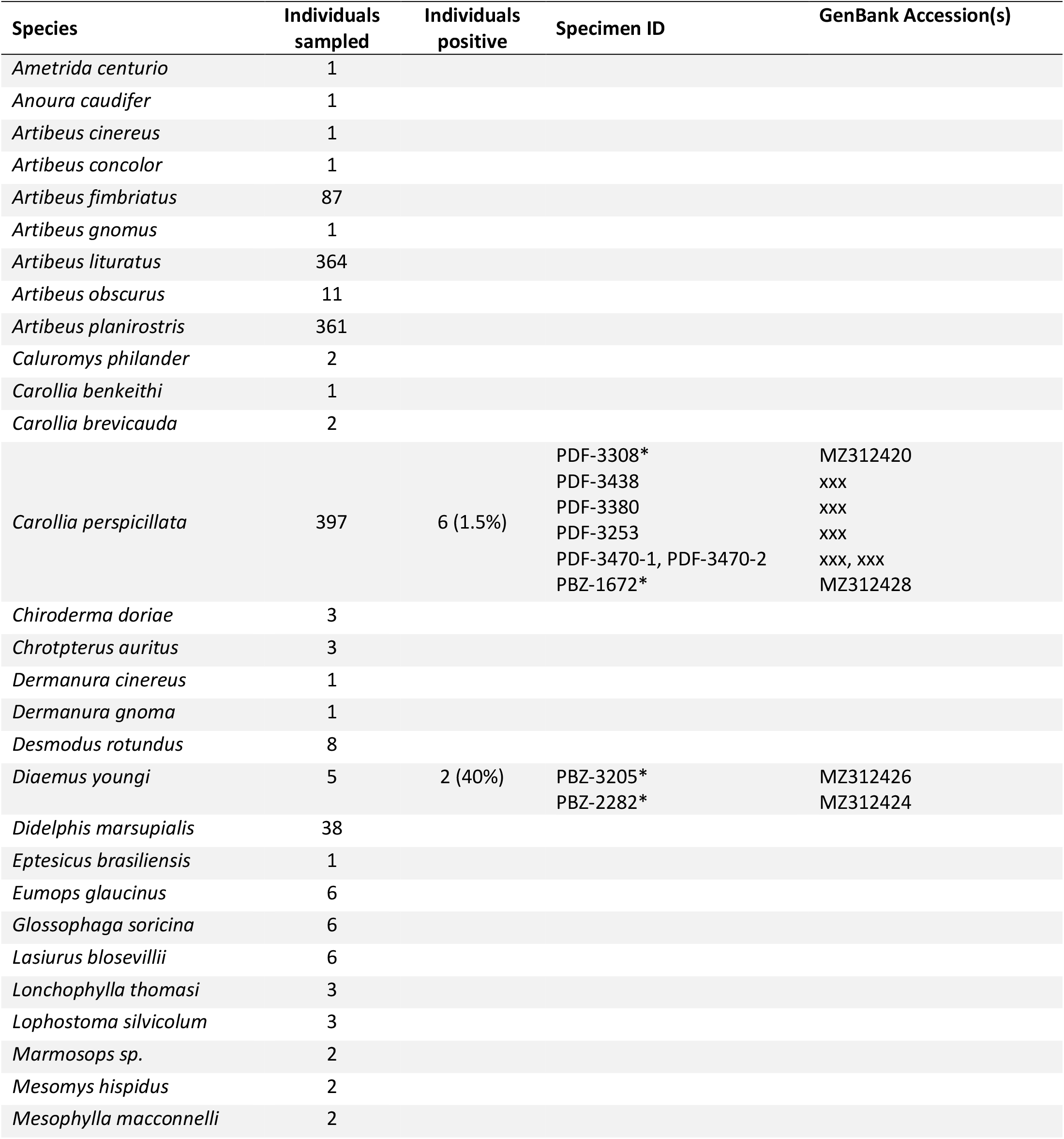
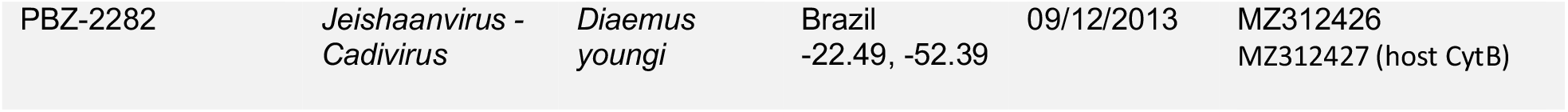
Full-genome viral sequences described in this study along with suggested taxonomic classification, host species, geographic origin (plus latitude/longitude where available), collection date, and associated GenBank accession numbers.

### Phylogenetics

To reconstruct the phylogenetic history of the seven novel genomes, we first collected from GenBank any orthoparamyxovirus sequence for which the full genome was available as well as those for which the majority of the polymerase (L) gene was available. We excluded the genus *Respirovirus* as none of our sequences fell within this group, but we included Sendai virus as an outgroup to root the phylogeny. Phylogenetic reconstruction was first performed using all available full- or nearly full-length L nucleotide sequences, which were aligned relative to their amino acid translations using MUSCLE (32 sequences and 8,048 bp, see Supplementary material). Bayesian maximum clade credibility trees were generated using BEAST v2.6.3 with no tip dating, BEAST model test averaging, a strict molecular clock, and a Yule model process prior. MCMC chains were run until trace plots demonstrated convergence and the estimated sample size (ESS) for all parameters was greater than 200 using Tracer v1.7.1. Phylogenetic trees were also built for each of the other genes individually (N, P, M, F, and RBP) using the same methods to compare topologies for any inconsistencies. Six sequences for which only the polymerase gene was available were excluded from these trees.

### Pairwise Amino Acid Sequence Comparison (PASC)

All orthoparamyxovirus sequences classified to the species level in GenBank for which any amount of L sequence was available were used for PASC analysis. This data consisted of several long sequences, but the majority of sequences in GenBank are very short fragments (∼500bp) generated by common PCR assays used for viral discovery [18] (Figure 1, right). Because these fragments are from assays targeting different regions and do not overlap, an alignment was created by manually. First, all full- or nearly full-length L sequences were aligned. Second, Tong-PanPMV (PCR assay targeting all paramyxoviruses) and Tong-RMH (PCR assay targeting sequences in *Respirovirus, Morbillivirus*, and *Henipavirus*) sequences were each aligned separately. Finally, an alignment with one full length L sequence and one sequence of either Tong-PanPMV or Tong-RMH was used as a reference with which to position the fragment region alignments joined onto the full L alignment backbone. Because there are no sequence gaps in the fragment regions of the alignments, this method is robust to its manual nature. The alignment is available as Supplementary material.

Pairwise distances between each sequence were generated using nucleotide percent identity or BLOSUM62 matrix scores and used to build histograms demonstrating the distributions of pairwise identities (PIDs) between viruses classified as the same species, as different species within the same genus, or as species within two different genera. This approach is in contrast to the NCBI-PASC tool, which uses a BLAST-based comparison calculation [21]. This analysis was performed for nucleotide as well as amino acid sequences. Where fragments did not overlap, no pairwise comparisons were calculated. For the short fragment histograms, only the corresponding regions were used within sequences for which full- or nearly full-length L was available. For amino acid alignments, percentages were calculated as the BLOSUM62 alignment score divided by the total possible score (i.e., 100% identity).

## Results

cPCR screening of samples collected in Brazil identified 15 novel paramyxovirus sequences from 6 bat species (*Carollia perspicillata, Diaemus youngi, Myotis riparius, Phyllostomus elongatus, P. hastatus*, and *Pteronotus parnellii*) (Supplementary Table 1). Based on the L gene phylogeny, two sequences, one from *P. hastatus* and one from *M. riparius*, clustered with the *Morbillivirus* genus (Supplementary Figure S1). The rest of the sequences clustered within an unnamed sister clade to *Jeilongvirus* and *Shaanvirus*, with the exception of one sequence from *P. parnellii*. This sequence clustered within the same group but was positioned on a single long branch (Supplementary Figure S1).

In Sabah, Malaysia, cPCR screening identified an additional 24 novel paramyxovirus sequences from 6 bat species (*Hipposideros cervinus, H. diadema, H. galeritus, Rhinolophus arcuatus, R. creaghi*, and *R. trifoliatus*) and one sequence from a moonrat (*Echinosorex gymnura*), an animal which is not closely related to rodents and is from the hedgehog family (Supplementary Table 2). All of these sequences clustered within either Clade 1 or Clade 2 of the *Shaanvirus* group in the L gene phylogeny (Supplementary Figure S1). The single sequence from *E. gymnura* is most closely related to Belerina virus, which was isolated from a European hedgehog (*Erinaceus europaeus*).

In total, 39 cPCR-positive samples were sent for high-throughput sequencing. Of these, we recovered seven novel genome sequences (Table 1). PDF-3137 and PBZ-1381 from Brazil belong to the *Morbillivirus* genus, clustering most closely with canine and phocine distemper viruses in the L gene phylogeny (Figure 2). PDF-3137 was found in a *Myotis riparius* bat in the family *Molossidae*, while PBZ-1381 was found in a *Phyllostomus hastatus* bat in the family *Phyllostomidae*. Both have the same genome arrangement as other morbilliviruses (Figure 3), and PDF-3137 has also been shown to utilize the morbillivirus receptors SLAMF1/NECTIN4 [22]. The single sequence from Sabah, Malaysia, PDF-0699, was found in a sample from a *Hipposideros galeritus* bat. This sequence was most closely related to the Clade 2 members of *Shaanvirus* and has the same two ATUs between the fusion and receptor binding proteins (Figures 2 and 3). The remaining four sequences from Brazil, PDF-3308, PBZ-1672, PBZ-3205, and PBZ-2282, comprise a monophyletic group with no conclusive phylogenetic placement into an existing genus (Figure 2). These four unclassified sequences were each characterized by the presence of a single ATU in the middle of the genome (Figure 3). A blastx search in NCBI of the ATU from each genome resulted in no hits to any other published sequence in GenBank. The ATUs of PDF-3308 and PBZ-1672 appear to be homologous and share 92% sequence identity, while PBZ-3205 and PBZ-2282 share <40% sequence identity with any of the other three sequences and only 25% identity to each other, suggesting they are not homologous. PDF-3308 and PBZ-1672 were found in *Carollia perspicillata*, while PBZ-3205 and PBZ-2282 were found in *Diaemus youngi*, both in the family *Phyllostomidae*.

**Figure 2.**
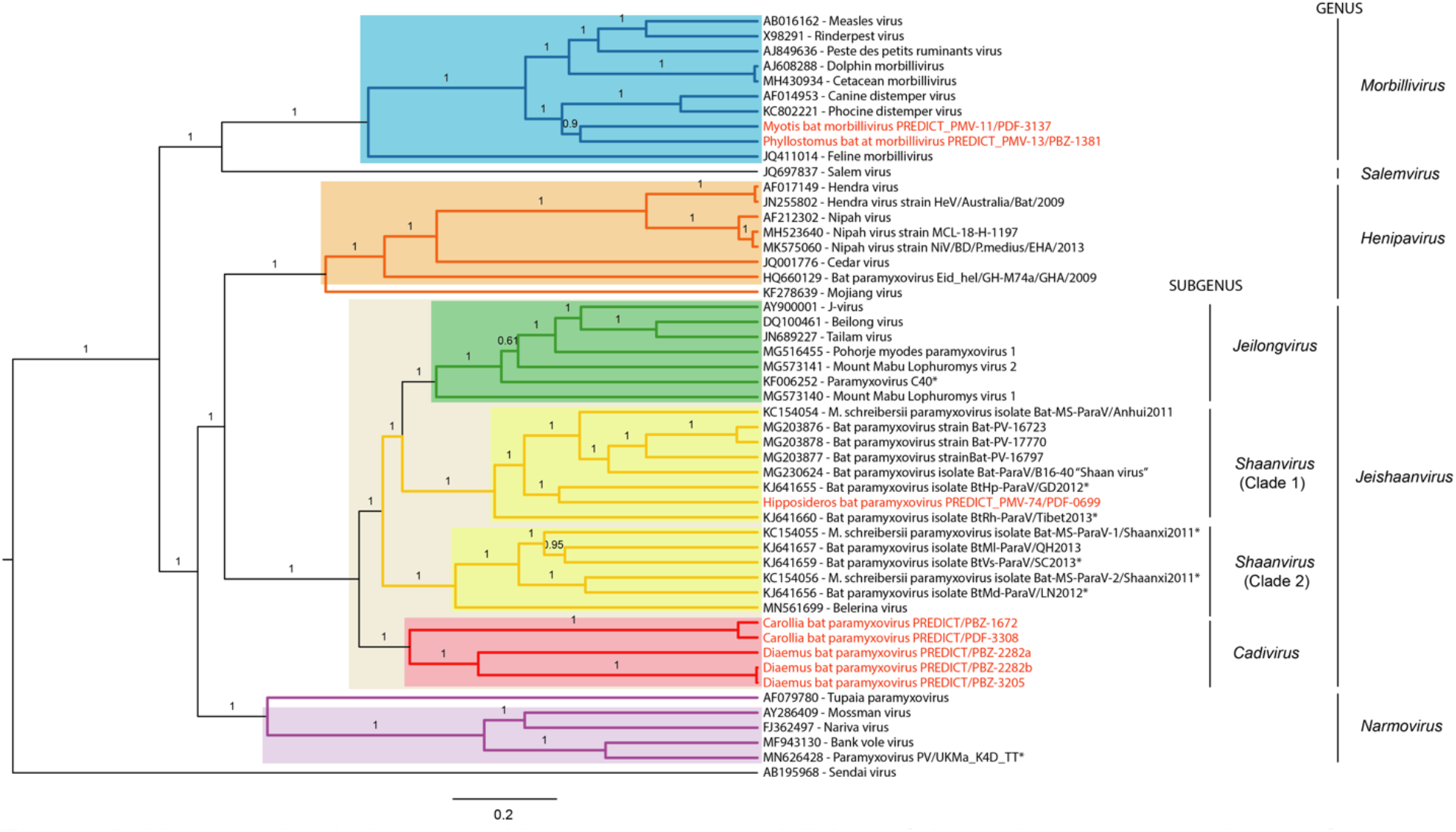
Nucleotide phylogeny with posterior probabilities of the orthoparamyxoviruses for which full- or nearly full-length L gene sequence is available. The genus *Respirovirus* is represented by a single sequence as outgroup, Sendai virus. Clade bars are colored by their current genus classifications which are also labeled on the right. Box highlights over clade bars represent taxonomic suggestions discussed here, including unclassifying Tupaia virus and Mojiang virus and formally naming the genus *Jeishaanvirus* including *Jeilongvirus, Shaanvirus* (Clades 1 and 2), and *Cadivirus* as subgenera.

**Figure 3.**
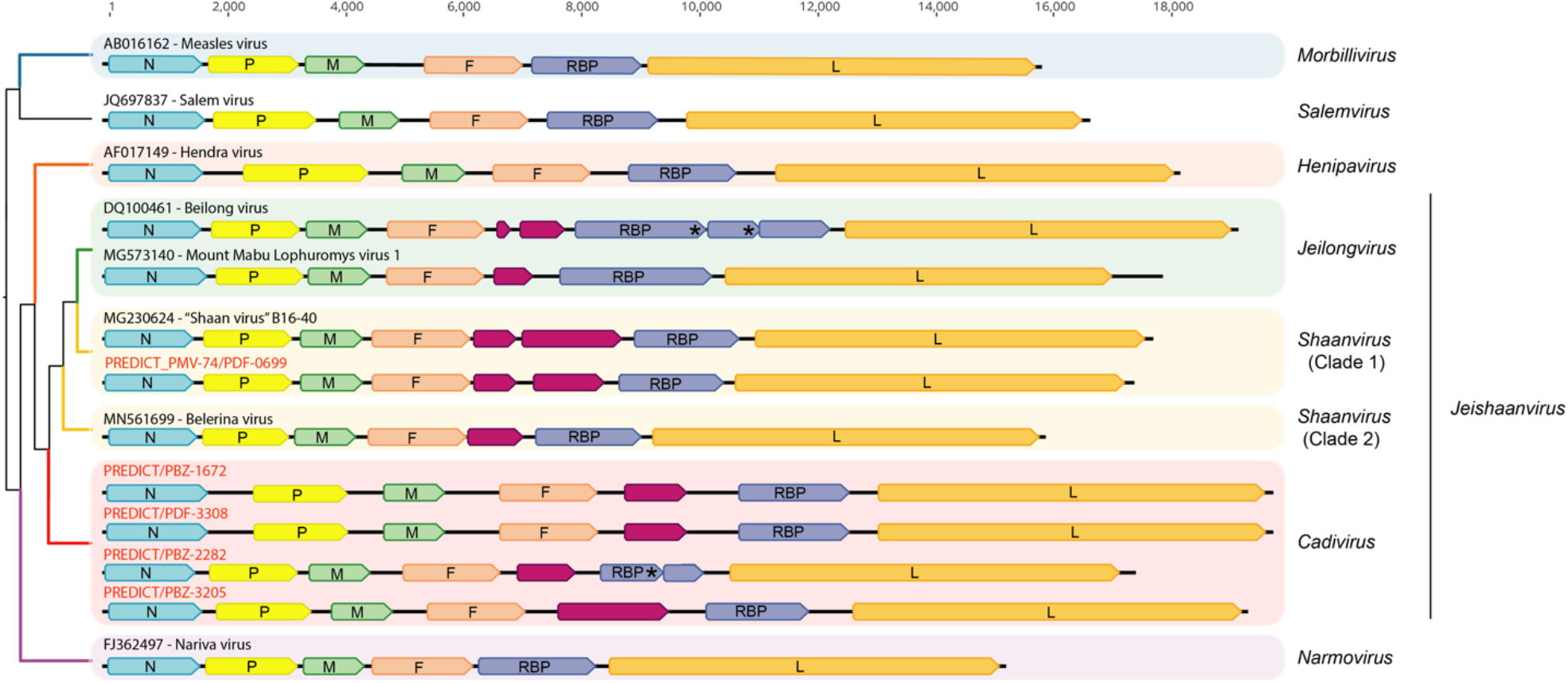
Genome organizations of representative sequences from each genus included in this study. Sequences are organized according to their L gene phylogeny, which is shown on the left. New genome sequences described here are labeled with red font, with the exception of the bat morbilliviruses (PDF-3137 and PBZ-1381) which share the same genome arrangement as all morbilliviruses. Where more than one genome arrangement is present in a single genus, a representative of each type is shown. All genome arrangements in the proposed new genus are shown. ORFs are shown by colored polygons, and the black lines are the entire length of the sequence. Both are illustrated to scale. ORFs are colored as follows: nucleocapsid (N gene): blue, phosphoprotein (P gene): yellow, matrix (M gene): green, fusion (F gene): peach, ATUs: magenta, receptor binding protein (RBP gene): purple, and polymerase (L gene): orange. Asterisks on RBP ORFs indicate premature stop codons.

We also examined the topology of the phylogenies constructed from the other five genes and found that these new sequences from Brazil consistently cluster as a monophyletic clade (Figure 4). During our investigations, we also noted that previously published sequences proposed as the new genus *Shaanvirus* form two clades which were not monophyletic in the L gene (Figure 2). In addition, these sequences were also paraphyletic in phylogenies constructed from all five other genes of the genome (Figure 4). Similarly, we found that Tupaia virus is not monophyletic with respect to the other *Narmovirus* sequences for the N, M, F, and RBP genes, where it resides independently on a single long branch (Figure 4).

**Figure 4.**
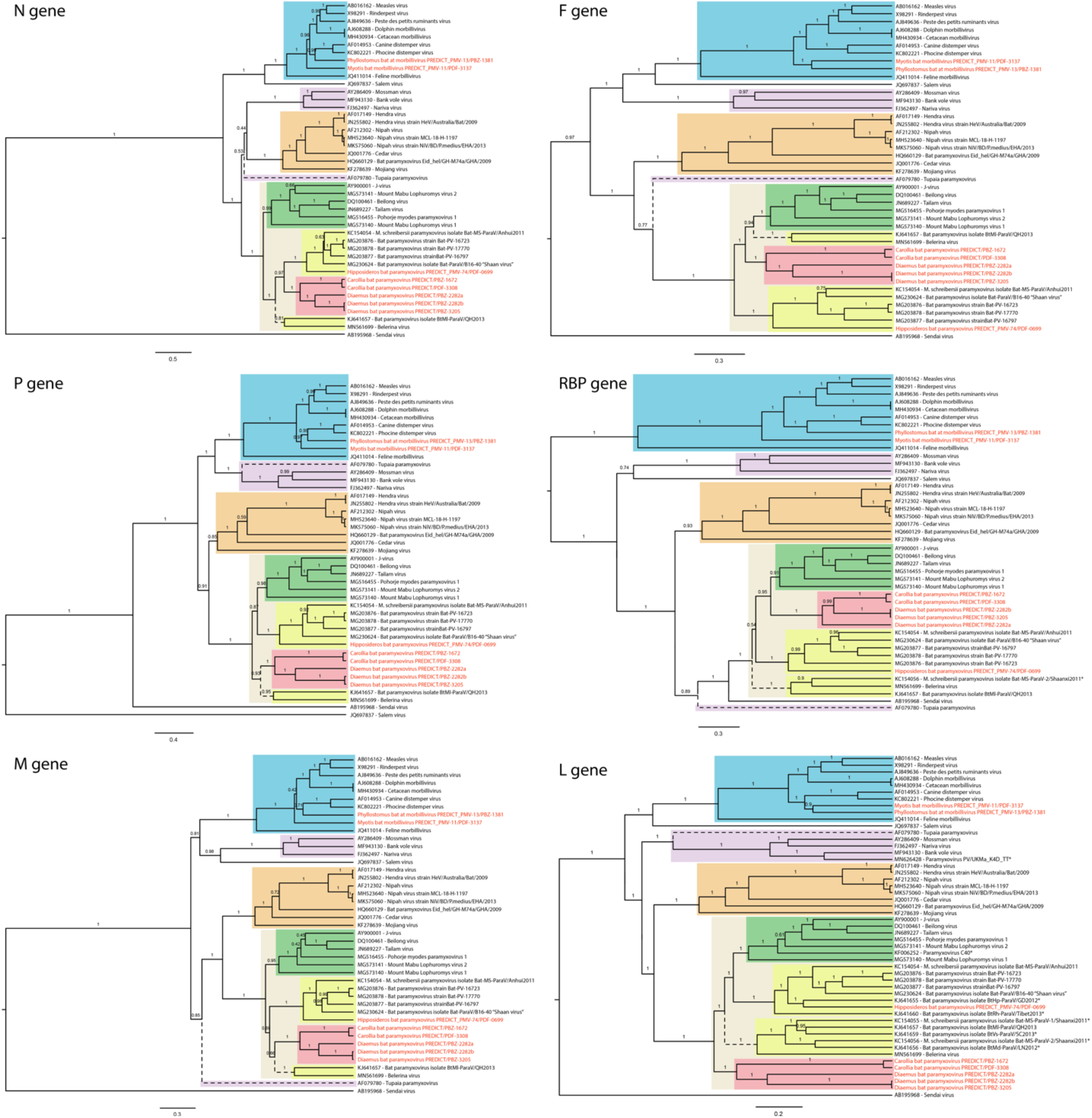
Nucleotide phylogenies for all genes, excluding ATUs. Clades are color-coded with the same scheme as Figure 1: *Morbillivirus*: blue, *Henipavirus*: orange, *Jeilongvirus*: green, *Shaanvirus*: yellow (both Clade 1 and Clade 2),*Cadivirus*: red, and *Narmovirus*: purple. Where sequences are incomplete, no sequence is shown in the gene tree. For clades that are not monophyletic by genus in one or more gene trees (i.e., paraphyletic *Shaanvirus* sequences and Tupaia virus), branches are shown as dotted lines. Shaded boxes represent current taxonomic classifications with the same color scheme as Figure 2.

To support classification of the four genomes from Brazil as a new genus, we implemented PASC at the nucleotide and amino acid levels. As no distinct cutoffs have been previously published for demarcation within *Paramyxoviridae* [4], we first performed PASC on all existing classified orthoparamyxoviruses within the ICTV-approved genera *Morbillivirus, Henipavirus, Jeilongvirus*, and *Narmovirus* to compare the distributions of PIDs for sequences of the same species, of the same genus, and of different genera (Figure 5). The distribution of pairwise amino acid identities (PAIDs) for sequences of the same viral species is well-separated from the distribution of PAIDs between different species above and below 96%; however, there is significant overlap in distributions of PAIDs between sequences of the same genus and sequences of different genera between 70 and 90% (Figure 5A). We suspected this may be due to the fact that many sequences submitted to GenBank are generated with the same commonly used consensus PCR assay, Tong-PanPMV and Tong-RMH. Since these assays each target a small, conserved region of the L gene, PIDs may be biased upwards when only this small fragment is available for comparison. Indeed, we found that when we looked at PAIDs only between sequences with 350 or more amino acids, the distributions became much more distinct (Figure 5B). We also generated histograms for each of the two consensus PCR assay fragments (Tong-PanPMV and Tong-RMH), which demonstrated that the average PAID between sequences of different genera shifted upwards from ∼65% to nearly 80%, eliminating the distinction observed when only long sequences are considered (Figure 5C and D).

**Figure 5.**
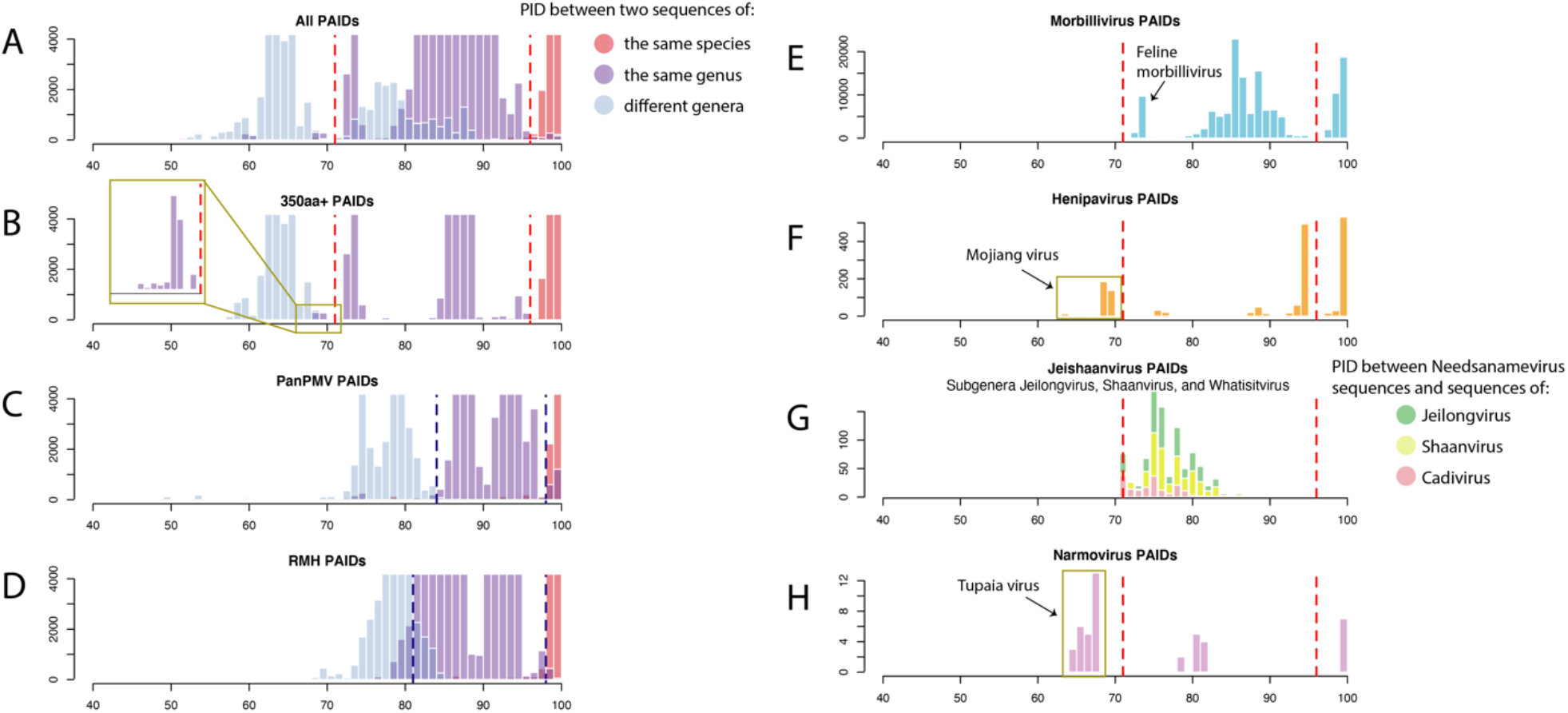
PASC histograms of pairwise amino acid comparisons of the L gene. Scores on the x-axis were calculated by scoring alignments with the BLOSUM62 matrix and dividing by the total possible score (100% identity). Y-axes indicate frequency. Gold boxes correspond to PIDs between sequences that do not conform to identified cutoff values based on their established classifications. Cutoff values used are shown as dotted lines, with red for full-length L sequences and blue for short fragments (Tong-PanPMV and Tong-RMH).

Although the histogram of PAIDs for sequences greater than 350aa is a significant improvement in demarcation, there is a small peak of PAIDs between sequences classified as belonging to the same genus that fall closer to the distribution of PAIDs between sequences classified as belonging to different genera (Figure 5B; gold box). To investigate this peak, we performed the same PASC analysis for each genus individually. The PAIDs of three of the classified genera (*Morbillivirus, Henipavirus*, and *Narmovirus*) mainly show distributions that would be expected for sequences of the same species or genus, but *Henipavirus* and *Narmovirus* each contain a small cluster of sequences distinctly less similar to the rest of the distribution (Figure 5F and H; gold boxes). We identified these PAIDs as *Henipavirus* sequences compared with Mojiang virus and *Narmovirus* sequences compared with Tupaia virus. A cluster of low similarity was also observed for *Morbillivirus* sequences compared with feline morbillivirus (Figure 5E). We decided to place a cutoff value at 71% for two sequences to belong to the same genus, as this results in all PAIDs between sequences from different genera falling below the cutoff and the majority of sequences from the same genus falling above this cutoff. This cutoff value supports the classification of Feline morbillivirus in the genus *Morbillivirus* but places Mojiang virus and Tupaia virus outside *Henipavirus* and *Narmovirus*, respectively.

Sequences classified in the genus *Jeilongvirus* all have PAIDs that fall above the cutoff. *Shaanvirus* sequences in both Clades 1 and 2 have PAIDs >73%, which would place the two in the same genus despite the fact that they are not monophyletic and have different genome arrangements. Further, PAIDs between *Shaanvirus* sequences, *Jeilongvirus* sequences, or the four novel sequences from Brazil compared to any other are all >71%, which would technically place each of these groups into a single genus despite vastly different biological characteristics (Figure 5G).

Identical analyses were performed at the nucleotide level, and the distributions of pairwise nucleotide identities (PNIDs) result in the same inferences as those of the PAIDs. The cutoffs for nucleotides were found to lie at approximately 55% and 80% for genus and species, respectively.

## Discussion

Relying only on genetic sequence and biological characteristics, we have provided evidence to support the classification of four novel paramyxovirus genomes (PDF-3308, PBZ-1672, PBZ-3205, and PBZ-2282) into a new subgenus, here putatively named *Cadivirus*. This classification is based on PASC analysis, which shows that these four sequences do not meet our defined cutoff criteria of 71% PAID or 55% PNID compared to *Shaanvirus* or *Jeilongvirus* sequences. We suggest that these three groups be classified into a single genus called *Jeishaanvirus*, with *Cadivirus, Shaanvirus* Clades 1 and 2, and *Jeilongvirus* as subgenera. Grouping them together is supported by the fact that these groups comprise the only orthoparamyxoviruses that have ATUs in the middle of the genome. However, the distinct characteristics of each, such as host taxa and geographic range, underline the need for further taxonomic distinction to the subgenus level. For example, jeilongviruses are exclusively found in rodents, while shaanviruses and cadiviruses are primarily found in bats, underscoring distinct evolutionary trajectories. A geographic range in South America is unique to the four novel sequences reported here, where no other complete genome paramyxoviruses from bats have been sequenced or classified to date. We additionally describe the first two published full-genome bat morbilliviruses from this geographic region (PDF-3137 and PBZ-1381). The finding that PDF-3137 shares the same receptor usage as all other morbilliviruses, SLAMF1 and NECTIN4, further supports its classification into the genus *Morbillivirus* [22](referred to as MBaMV). The only other known bat morbillivirus sequences were also collected from Phyllostomid bats in Brazil but were not completely sequenced [2].

We also show that our single sequence from Sabah, Malaysia, PDF-0699, shares PIDs consistent with *Shaanvirus* Clade 2 sequences and has the same arrangement of ATUs in the genome as other members of this clade. Only two Clade 2 members have published full genomes (Belerina virus and QH2013) and each has only one ATU that is not homologous to either of those present in the first clade. The two *Shaanvirus* clades are also consistently paraphyletic in every other gene of the genome (Figure 4). While the PASC analysis PAIDs and PNIDs of the *Shaanvirus* sequences from both clades do not show conclusive evidence that they should be considered separate genera, the lack of monophyly of every gene in the genome and the presence of a different ATU provide strong evidence that they should be classified separately. We suggest that the sequences in the monophyletic clade clustering with the virus originally named “Shaan virus” (B16-40) and including PDF-0699 be classified in the subgenus *Shaanvirus*. Those in the paraphyletic clade should be placed in a separate subgenus.

In performing our analysis to determine PASC cutoff values, we found that pairwise comparisons of sequences within the same genus to Mojiang virus and Tupaia virus generated values that fell below our same-genus cutoff values (Figure 5). This finding raises a question as to whether these sequences should actually be considered as members of the genera in which they are currently classified. PASC is just one tool that can be used to taxonomically classify these sequences, and other characteristics should be considered to determine if a sequence should be classified within an existing genus, such as receptor usage, host range, and presence of ATUs (Table 2). For example, feline morbillivirus PAIDs fall extremely close to the 71% cutoff, but this virus is highly consistent with other morbilliviruses in other contexts. The morbilliviruses have a broad range among mammals, use SLAM1/NECTIN4 cell receptors, and have no ATUs – all of which are consistent with feline morbillivirus, which also clusters with the monophyletic morbillivirus clade in every gene. The same does not apply to Tupaia virus and the other members of *Narmovirus*; however, their cell receptor usages are currently not known. Tupaia does not cluster within the *Narmovirus* clade in the N, M, F, and RBP genes and falls below the 70% PAID cutoff by PASC analysis. Mojiang virus also shows significant deviation from the traits of other *Henipaviruses*. Mojiang virus is rodent-borne and is unable to use ENFB2/3 as a receptor, whereas other members of this genus are bat-borne and use ENFB2/3. Mojiang virus is consistently monophyletic with other henipaviruses, but PASC analysis also places this sequence below the 70% PAID cutoff. While Feline morbillivirus characteristics are in line with other morbilliviruses despite its lower PIDs, the same is not true for Mojiang virus and Tupaia virus, which should have their current genus classifications reconsidered.

**Table 2.** Genera of the paramyxovirus subfamily *Orthoparamyxovirinae*, with the exception of *Salemvirus* and *Ferlavirus* which have only one classified member and *Respirovirus* and *Aquaparamyxovirus* which are outside the scope of this study. For each genus, the host taxa, host cellular receptor (if known), and the number of ATUs are shown. In addition, if any classified member of that genus represents an exception to those traits, it is listed along with the differing trait(s) in the last column.

| Genus | Host | Receptor | ATUs | Exceptions |
| --- | --- | --- | --- | --- |
| *Morbillivirus* | mammals | SLAMF1/NECTIN4 | none |  |
| *Henipavirus* | bats | ENFB2/3 | none | Mojiang virus: host is a rodent, does not use ENFB2/3, falls below PASC cutoff |
| *Jeilongvirus* | rodents | unknown | two |  |
| *Shaanvirus*<br>(Clade 1) | bats | unknown | two |  |
| *Shaanvirus*<br>(Clade 2) | bats | unknown | one | Belerina virus: host is a hedgehog |
| *Cadivirus* | bats | unknown | one |  |
| *Narmovirus* | rodents | unknown | none | Tupaia virus: does not share monophyly, falls below PASC cutoff |

Pairwise amino acid sequence comparison (PASC) analysis has previously been suggested as a tool to rapidly classify viruses when only the genetic sequence is available [21]. When distributions of pairwise identities (PIDs) form distinct distributions separable by a single cutoff value, this analysis can be very effective. With paramyxoviruses, however, no such cutoffs have yet been determined and considerable difficulty has been highlighted in attempts to do so [4]. Here, we perform PASC analysis separately from the NCBI platform and used only the conserved polymerase gene. Using all available sequences in GenBank, we found considerable overlap in distributions of PIDs between sequences of the same genus or of different genera, with no single cutoff value appearing appropriate for delineating between the two, which is in agreement with the NCBI-PASC tool. However, we found that by separating PIDs by the length of the sequences being compared, much clearer cutoff values can be attained. Because the small sequence fragments prevalent in GenBank are generated using a consensus PCR assay that targets a conserved region of the genome, the PID comparisons using these sequences are biased upwards considerably. By restricting the sequences used to determine cutoff values to those with more than 350aa (or 1000bp), we demonstrate complete separation between sequence comparisons of the same genus and between different genera with a considerable gap in between, significantly reducing the ambiguity in cutoff placement. When considering PIDs between sequences shorter than this length, our cutoff values were shifted upwards by 4-6% and have a much less generous distinction between same or different genera, but may still be useful for classification where PIDs for the sequence in question are not near these cutoff values. One of the most important findings of this study is that short fragment sequences in GenBank should not be considered in the NCBI PASC analysis tool.

Taxonomic classification, particularly when it comes to viruses, is a task that requires continuous reevaluation and flexibility to keep up with new information when it becomes available. Previous classifications within the family *Paramyxoviridae* have changed several times in recent years with the discovery of new sequences, and the suggestions outlined here will undoubtedly come under review with the addition of new information in the future. The taxonomic changes described here are presented for discussion, but have not been endorsed by the ICTV Executive Committee and may ultimately differ from those which are approved. The avalanche of new sequences and information emerging for paramyxoviruses each year sets a pace that can quickly negate our best classification efforts. Future research should focus on determining the functions of the unknown ATUs in many of these genomes and on receptor usage for those which it is yet unknown, which will help illuminate borderline cases. Should researchers wish to embark on classification schemes using only PASC, careful consideration should be made to ensure that cutoff values being used are appropriate for the length of the sequences being considered. This is highlighted by our finding that PID distributions generated by short or long sequence lengths are considerably inconsistent. Finally, we emphasize the critical importance of the generation of full genomes where possible when describing novel viruses from wildlife that have been first identified using cPCR. In cases where genome sequencing is successful, the additional information that can be gleaned from the genome is not only highly valuable for classification of new sequences, but also for refinement of existing taxonomic classifications.

## Acknowledgments

We wish to thank the PREDICT Consortium, all of whom have contributed to the development of protocols and the curation of field sampling and data collection over the last ten years. We would also like to thank the governments and our partners in Brazil and in Sabah, Malaysia.

---

**Supplementary Figure S1.**
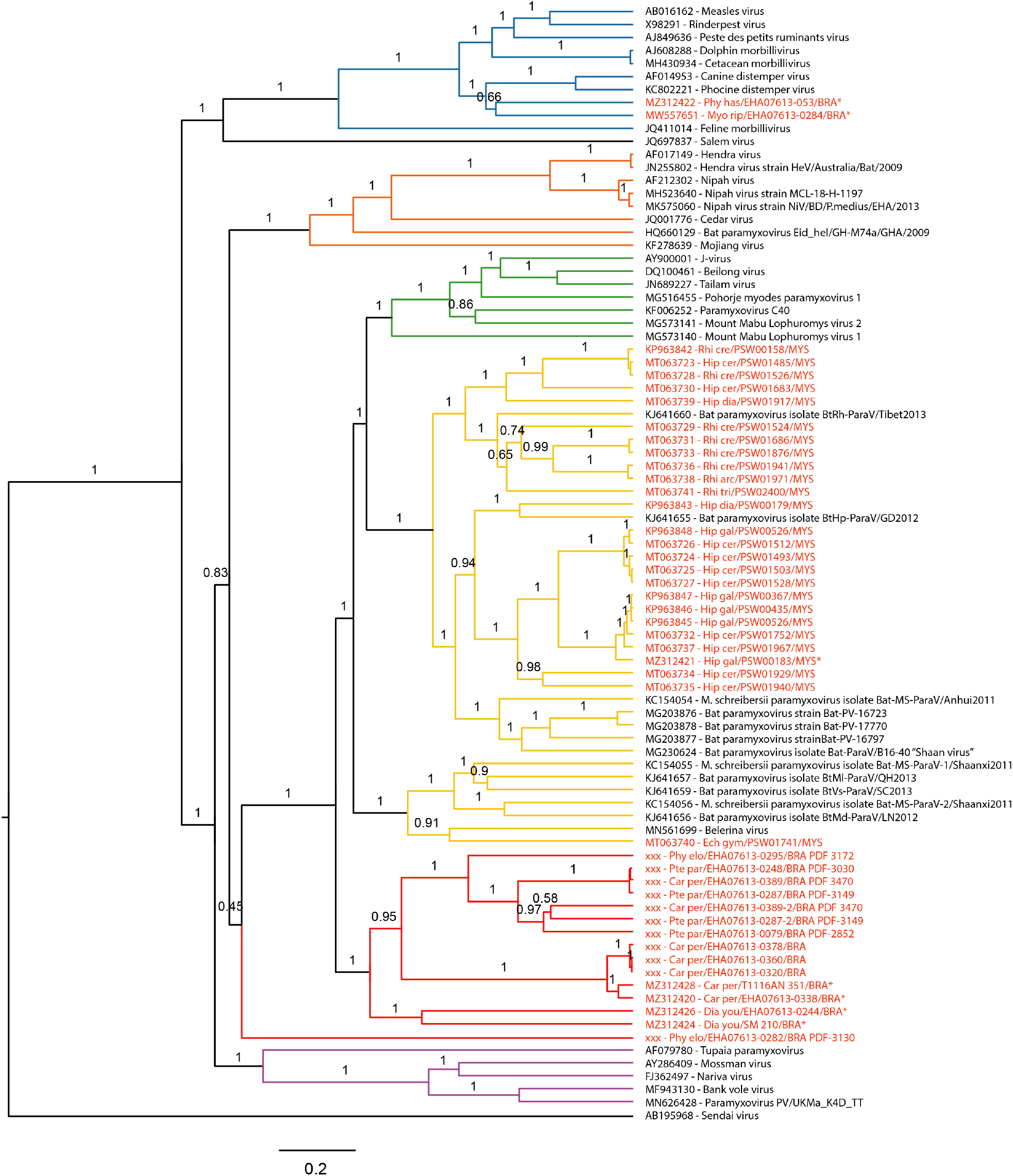
Phylogeny of all PCR fragments sequenced as part of this study with other classified paramyxoviruses. Names highlighted in red are those sequenced in this study, and those denoted with an asterisk are those for which full genomes were recovered. Clade bar colors are consistent with the taxonomic classifications in Figure 1. Some posterior probabilities near the tips of the tree have been removed for clarity.

**Supplementary Table 1.**
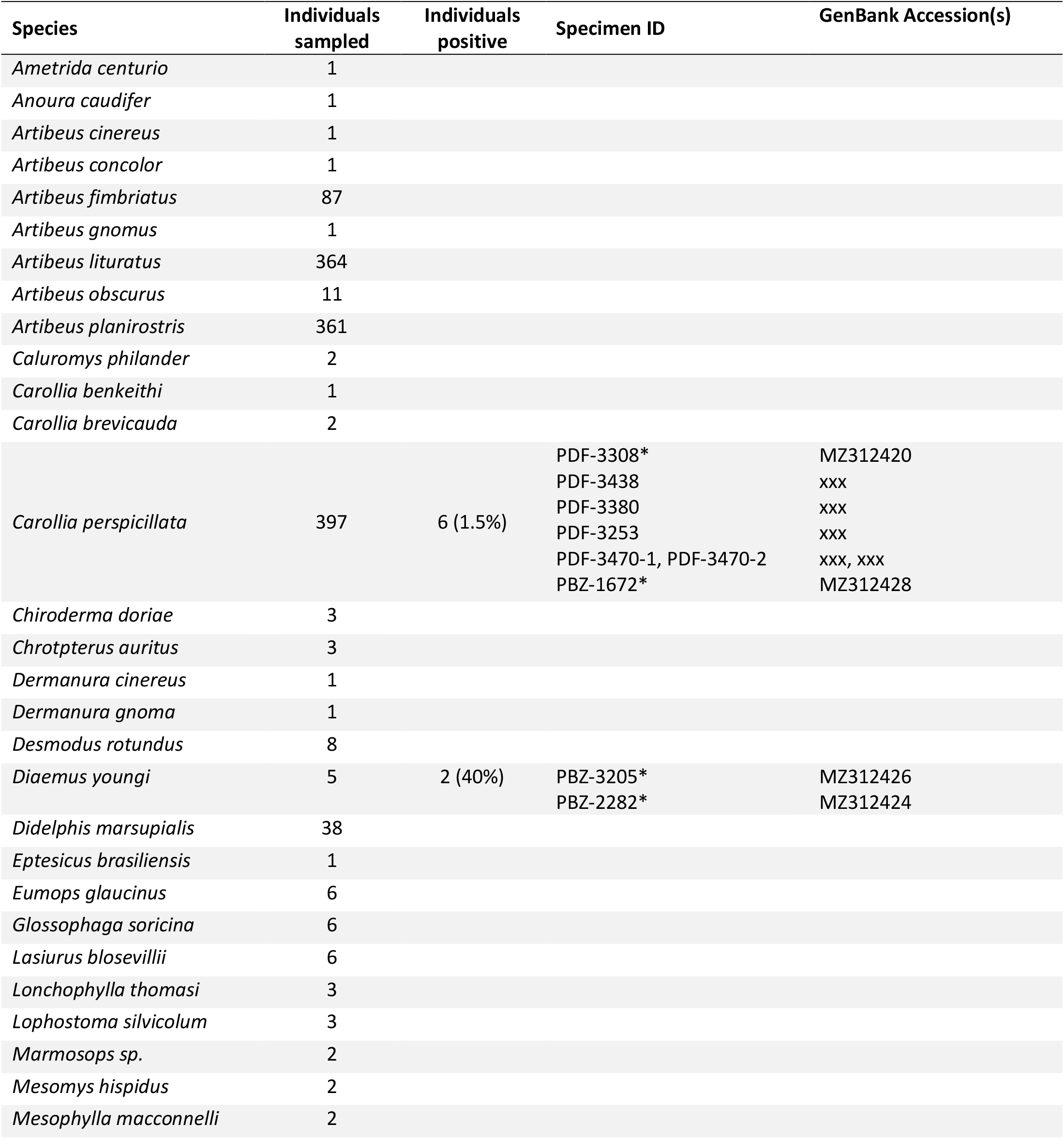
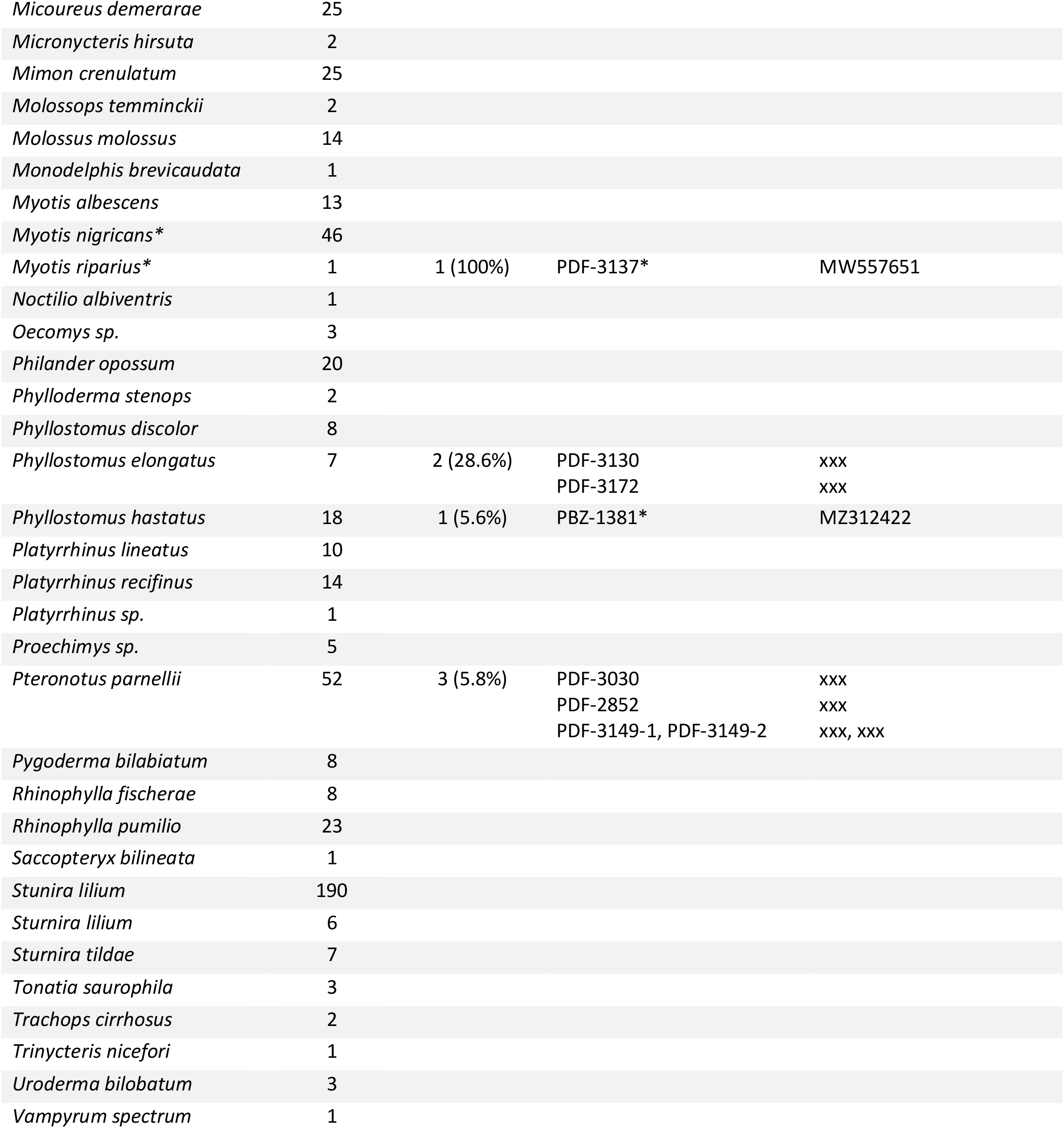
List of species sampled in Brazil, number of individuals tested from each species, and number of individuals found to be positive for paramyxoviruses. Percentages in parentheses are the positivity rate for that species. Names of viruses identified and corresponding GenBank accessions are also shown. Note that multiple virus names and GenBank accessions shown on the same line indicate coinfection of a single individual.

**Supplementary Table 2.**
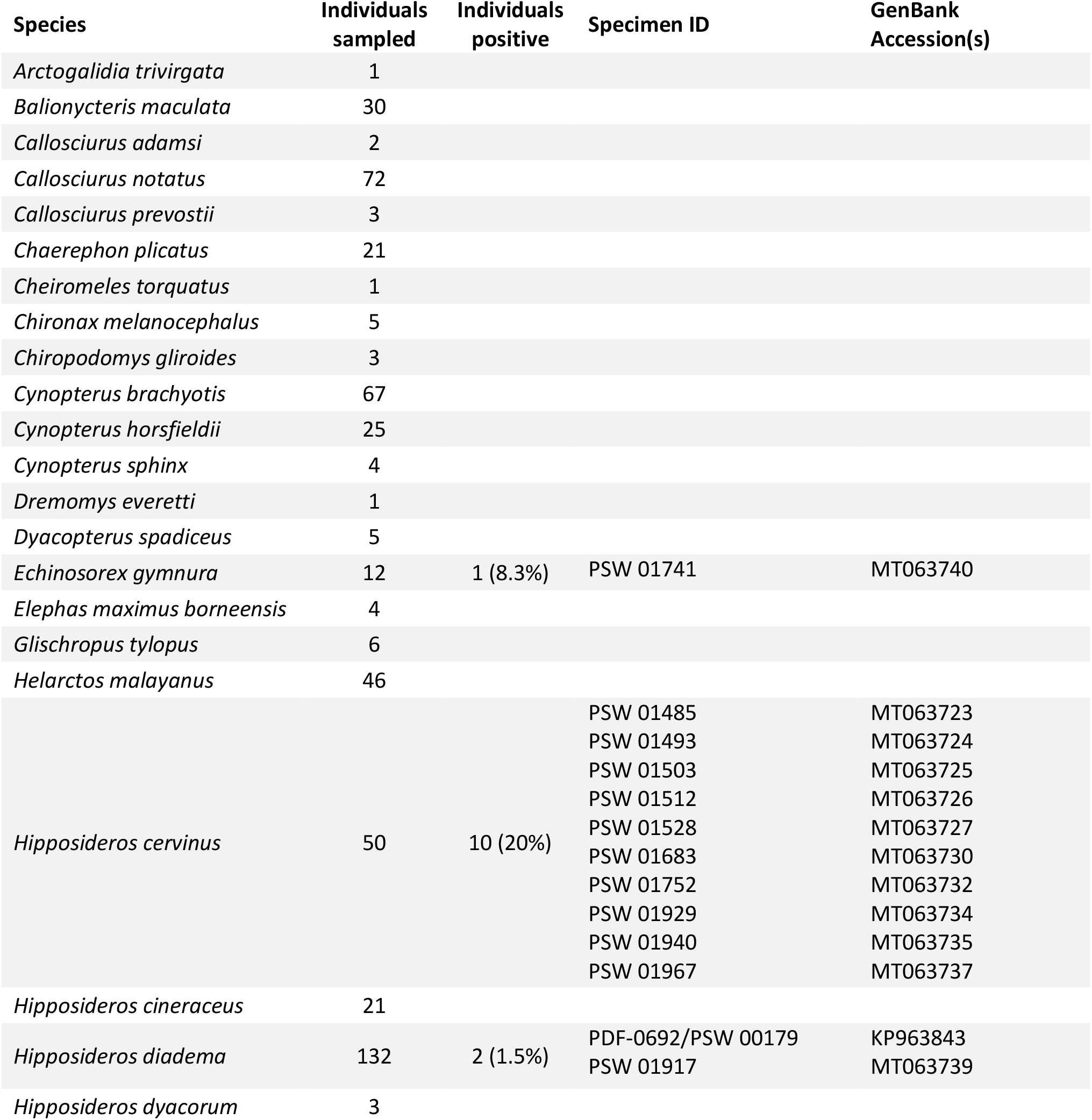
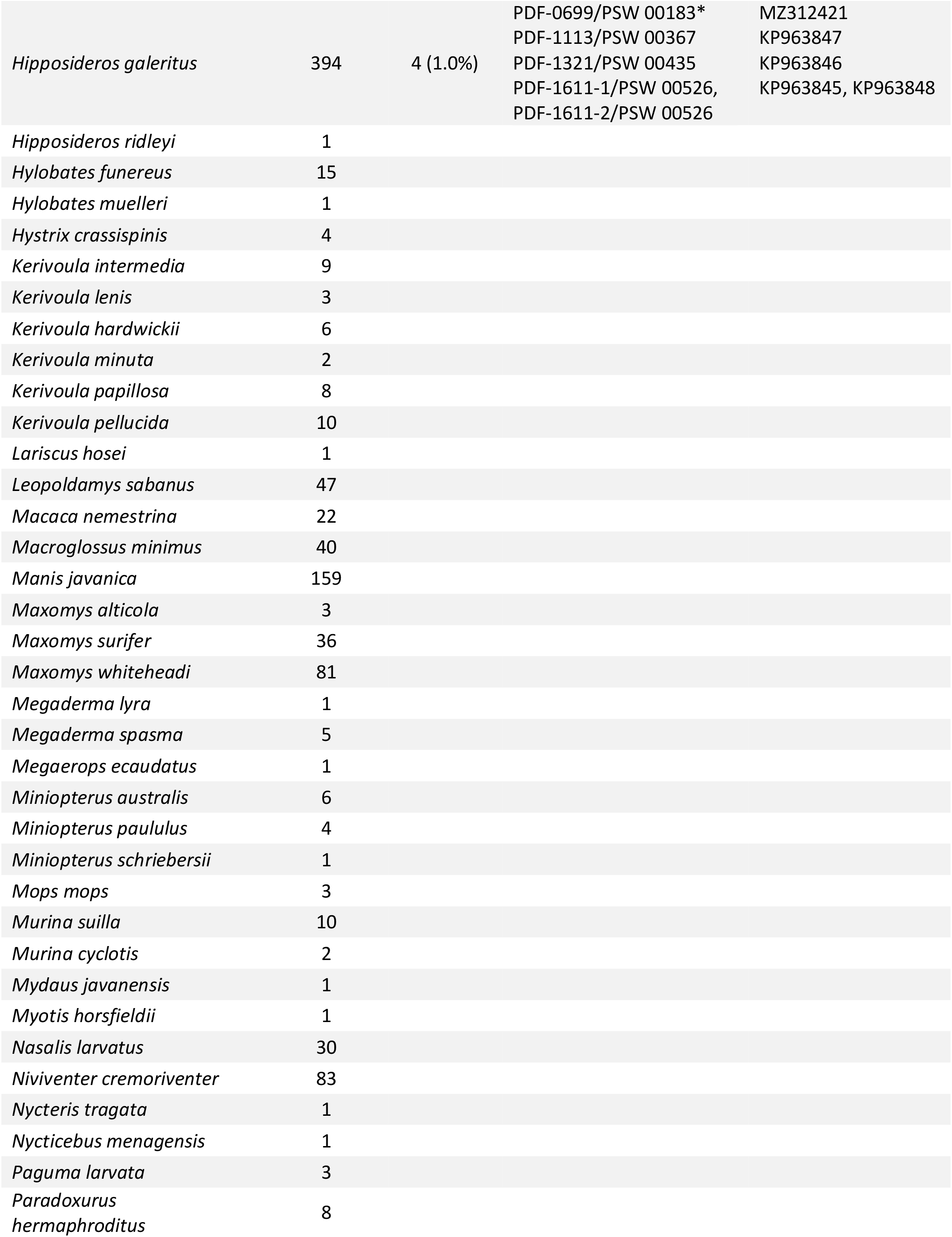
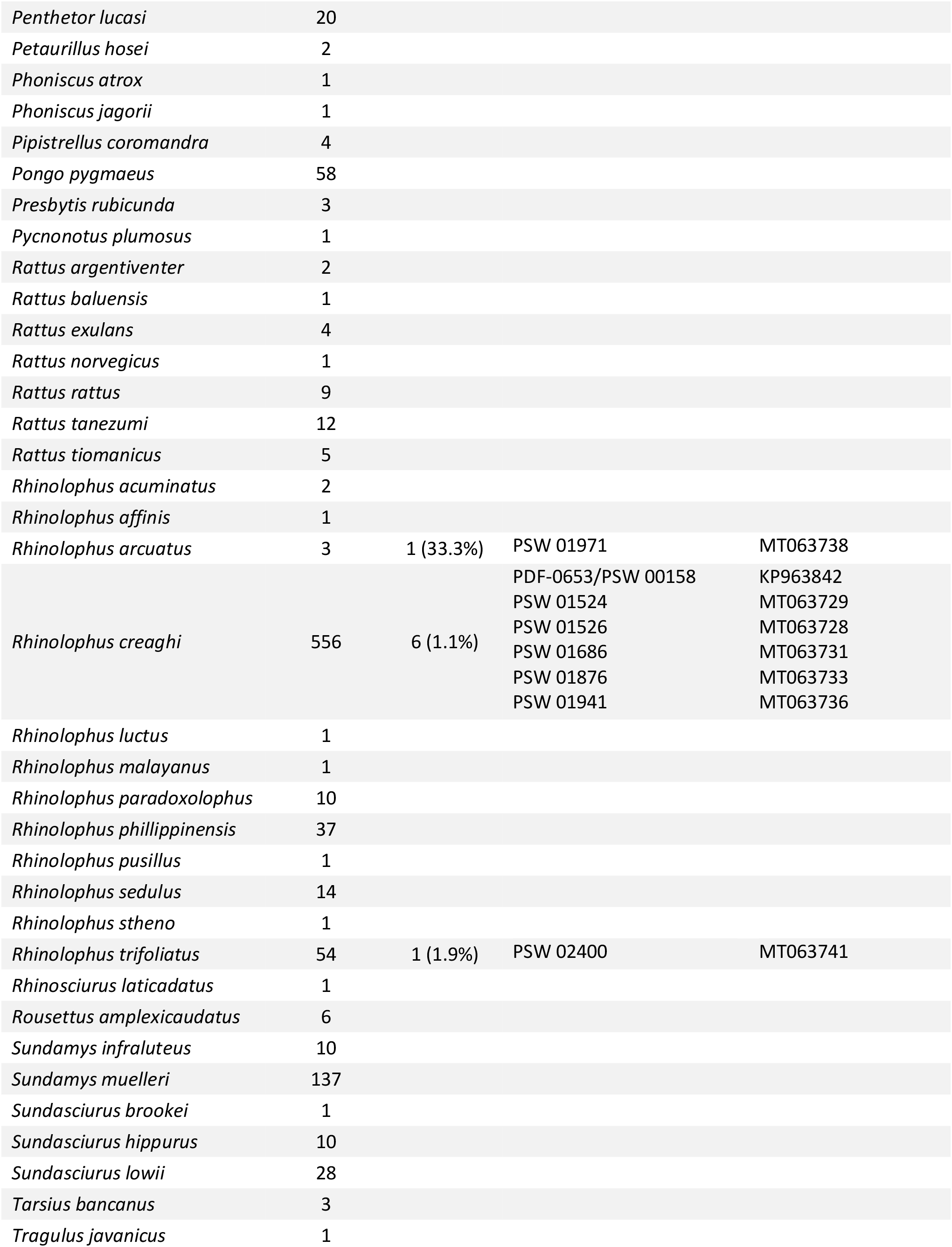
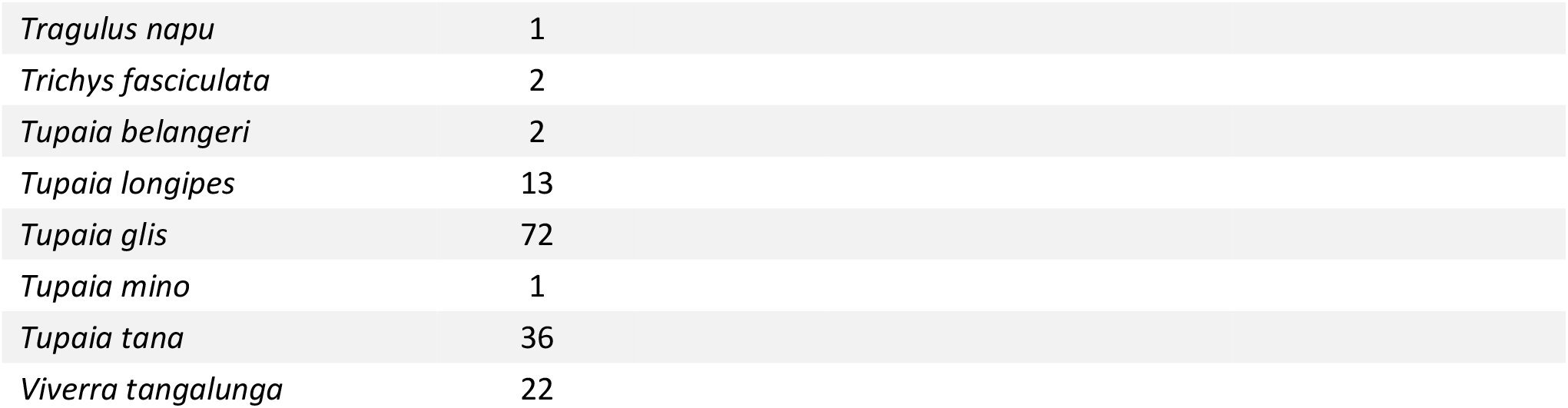
List of species sampled in Sabah, Malaysia, number of individuals tested from each species, and number of individuals found to be positive for paramyxoviruses. Percentages in parentheses are the positivity rate for that species. Names of viruses identified and corresponding GenBank accessions are also shown. Note that multiple virus names and GenBank accessions shown on the same line indicate coinfection of a single individual. Viruses marked with an asterisk are those for which full genome sequence was recovered.

**Supplementary Table 3.** Accession numbers in NCBI Sequence Read Archive (SRA) of fastq files obtained from sequenced samples.

| **Virus** | **Accession** |
| --- | --- |
| PDF-3137 | TBD |
| PBZ-1381 | TBD |
| PDF-0699 | TBD |
| PDF-3308 | TBD |
| PBZ-1672 | TBD |
| PBZ-3205 | TBD |
| PBZ-2282 | TBD |

